# Blocking abundant RNA transcripts by high-affinity oligonucleotides during transcriptome library preparation

**DOI:** 10.1101/2022.03.11.483910

**Authors:** Celine Everaert, Jasper Verwilt, Kimberly Verniers, Niels Vandamme, Alvaro Marcos Rubio, Jo Vandesompele, Pieter Mestdagh

**Author notes:** “The authors wish it to be known that, in their opinion, the first two authors should be regarded as joint First Authors”. ”The authors wish it to be known that, in their opinion, the last two authors should be regarded as joint Last Authors”.

## Abstract

RNA sequencing has become the gold standard for transcriptome analysis but has an inherent limitation of challenging quantification of low-abundant transcripts. In contrast to microarray technology, RNA sequencing reads are proportionally divided in function of transcript abundance. Therefore, low-abundant RNAs compete against highly abundant - and sometimes non-informative - RNA species. We developed an easy-to-use strategy based on high-affinity RNA-binding oligonucleotides to block reverse transcription and PCR amplification of specific RNA transcripts, thereby substantially reducing their abundance in the final sequencing library. To demonstrate the broad application potential of our method, we applied it to different transcripts and library preparation strategies, including YRNAs in small RNA sequencing of human blood plasma, mitochondrial rRNAs in both 3’ end sequencing and long-read sequencing, and MALAT1 in single-cell 3’ end sequencing. We demonstrate that the blocking strategy is highly efficient, reproducible, specific, and generally results in better transcriptome coverage and complexity. Furthermore, our method does not require modifications of the library preparation procedure apart from simply adding blocking oligonucleotides to the RT reaction and can thus be easily integrated into virtually any RNA sequencing library preparation protocol.

## INTRODUCTION

RNA sequencing has become the gold standard for transcriptome characterization. Numerous RNA sequencing library preparation procedures have been developed to quantify various RNA biotypes, including amongst others polyA+ RNA sequencing, total RNA sequencing, 3’ end RNA sequencing and small RNA sequencing. Regardless of the library preparation method, RNA sequencing reads are distributed across RNA transcripts proportionally to their abundance. Consequently, highly abundant RNA species, often deemed non-informative, can dominate the RNA sequencing library and hamper the detection of lower abundant transcripts. A well-known example is ribosomal RNA (rRNA), which typically accounts for more than 80% of all RNA transcripts (O’Neil et al., 2013b) in cellular or tissue RNA. Another well-documented example is RNY4 fragments. YRNAs are non-coding, evolutionary conserved RNA species with a length of 80-110 nucleotides. Four human YRNAs are known: hY1, hY3, hY4, and hY5 (Hendrick et al., 1981). YRNAs are readily fragmented in cells undergoing apoptosis in a caspase-dependent, Dicer-independent manner (Nicolas et al., 2012; Rutjes et al., 1999). The resulting fragments reside in cultured cells (Nicolas et al., 2012), solid tumors (Meiri et al., 2010), and multiple biofluids (Dhahbi et al., 2013; Ishikawa et al., 2017; Ninomiya et al., 2015). More specifically, a 30-33 nucleotide 5’-end hY4 fragment is abundantly present in human blood plasma, serum, and saliva potentially serving a physiological function (Ninomiya et al., 2019). In small RNA sequencing libraries of serum or plasma RNA, this fragment can account for more than 30% of all reads (Dhahbi et al., 2014; Yan et al., 2017) and even up to 70% in platelet-rich blood plasma. Superfluous amounts of hY4 fragments negatively impact the library complexity, requiring deeper sequencing to retrieve information about the other small RNA species in the library.

Removing sequence fragments derived from these excessively abundant transcripts from a sequencing library is instrumental in obtaining sufficient coverage of the informative fraction of the transcriptome without having to sequence libraries to extreme depth with diminishing returns. Several workarounds have been proposed to tackle this problem. The concentration of rRNA is reduced using many different strategies: subtractive pull-down (O’Neil et al., 2013a; Pang et al., 2004; Stewart et al., 2010; Su and Sordillo, 1998) (as in the old Ribo-Zero Gold Kit, the Ambion MICROBExpress Bacterial mRNA Enrichment kit and the Life Technologies RiboMinus Transcriptome Isolation Kit), gel excision (McGrath et al., 2008), probe-directed RNase H digestion (Benes et al., 2011; Huang et al., 2020; Morlan et al., 2012) (as in the new Ribo-Zero Gold Kit), Cas9-directed cDNA digestion (also named DASH) (Gu et al., 2016; Prezza et al., 2020), not-so-random primers (Armour et al., 2009; Arnaud et al., 2016), duplex-specific nuclease (DSN) depletion (Bogdanova et al., 2009; Yi et al., 2011), Probe-Directed Degradation (PDD) (Archer et al., 2015, 2014), rRNA poly(A) clipping (Naarmann-de Vries et al., 2022) and EMBR-seq (Wangsanuwat et al., 2020). These methods can in principle be applied for any other unwanted sequence. Instead of removing abundant transcripts, specific transcripts of interest can also be enriched using biotinylated probes, magnetic bead-linked probes, or capture arrays (Briese et al., 2015; Clark et al., 2015; Levin et al., 2009; Mercer et al., 2014, 2011; Morlion et al., 2021). Alternatively, methods like 3’ end sequencing apply poly(A)-priming to convert polyadenylated RNAs to cDNA for further library preparation. Several studies have compared the performance of some of these depletion methods and pointed towards discrepancies in efficiency and specificity (Bhagwat et al., 2014; Herbert et al., 2018a; Petrova et al., 2017; Zhao et al., 2014).

Methods developed for hybridization based small RNA depletion are often labor-intensive and result in loss of material by washing steps (Van Goethem et al., 2016). CRISPR-based technologies generally include PCR and multiple washes (Hardigan et al., 2019), making the protocol significantly longer and more prone to material loss. Likewise, pull-down methods also require several washing steps and tend to perform inconsistently (Petrova et al., 2017). Additionally, their efficiency drops significantly when applied to fragmented RNA in e.g., biofluids or formalin fixed tissues (Herbert et al., 2018b). All current technologies require the implementation of multiple steps and substantially increase the hands-on time and compromise the repeatability of the library preparation.

Researchers frequently use oligonucleotides containing modified nucleic acids due to their increased melting temperature, high binding specificity, or stability (Duffy et al., 2020). One example is locked nucleic acid (LNA), which contains an oxymethylene bridge between the 2’ oxygen and 4’ carbon molecules. This “locked” structure provides higher affinity and mismatch discrimination. Because of this, LNAs have been used in multiple applications (Breitenbuecher et al., 2014; Singh et al., 1998; Zhang et al., 2012). Interestingly, LNA oligonucleotides have been used to block the PCR amplification of unspliced transcripts (by targeting the intronic sequence) (Hummelshoj et al., 2005) or wild-type transcripts when the mutated version is of interest (Dominguez and Kolodney, 2005; Oldenburg et al., 2008; Vliegen et al., 2015). A patent describing the use of LNA oligonucleotides to block reverse transcription and amplification of hemoglobin mRNA from whole blood during RT-qPCR (Russell et al., 2006) further exemplifies their potential and applicability for depletion purposes. However, this idea has never been extensively evaluated and applied to massively parallel sequencing techniques, such as Oxford Nanopore Sequencing or single-cell RNA sequencing. Importantly, implementation of an LNA-based reverse transcription blocking step would require only one extra pipetting step during library preparation.

Here, we describe an easy-to-implement method using LNA-modified oligonucleotides that bind unwanted RNA transcripts and block their reverse transcription and PCR amplification during RNA sequencing library preparation. We applied our method to different abundant RNA species and RNA sequencing library preparation strategies, including small RNA sequencing, 3’ end sequencing, long-read sequencing, and single-cell 3’ end sequencing. We demonstrate that the applied method, which requires only one additional step in the library prep procedure, is highly efficient and does not affect quantification of untargeted genes.

## MATERIAL AND METHODS

### YRNA blocking in human blood plasma samples

#### Samples and sample collection

For the healthy donor experiments, we drew venous blood from an elbow vein of two healthy donors in three EDTA tubes (BD Vacutainer Hemogard Closure Plastic K2-Edta Tube, 10 ml, #367525) using the BD Vacutainer Push blood collection set (21G needle). We collected the blood samples according to the Ethical Committee of Ghent University Hospital approval EC/2017/1207, following the ICH Good Clinical Practice rules, and obtained written informed consents from all donors. We inverted the tubes 5 times and centrifuged within 15 minutes after blood draw (400 g, 20 minutes, room temperature, without brake). Per donor, we pipetted the upper plasma fraction (leaving approximately 0.5 cm plasma above the buffy coat) and pooled in a 15 ml tube. After gently inverting, five aliquots of 220 μl platelet-rich plasma (PRP) were snap-frozen in 1.5 ml LoBind tubes (Eppendorf Protein LoBind microcentrifuge tubes Z666548 - DNA/RNA) in liquid nitrogen and stored at −80 °C. We centrifuged the remaining plasma (800 g, 10 minutes, room temperature, without brake) and transferred to a new 15 ml tube, leaving approximately 0.5 cm plasma above the separation. Next, we centrifuged this plasma a 3^rd^ time (2500 g, 15 minutes, room temperature, without brake), and transferred it to a 15 ml tube, leaving approximately 0.5 cm above the separation. The resulting platelet-free plasma (PFP) was gently inverted, snap-frozen in five aliquots of 220 μl and stored at −80 °C. The entire plasma preparation protocol took less than two hours. We isolated RNA from 200 μl PRP or PFP. For the spike-in RNA titration experiment, the protocol was identical, except for the fact 4 EDTA tubes of 10 ml were used and that the second centrifugation step was different (1500 g, 15 minutes, room temperature, without brake).

For the cancer patient experiment, plasma samples are acquired from ProteoGenex (Inglewood, United States of America) under EC/2017/1515 from Ghent University Hospital. Blood was collected in EDTA vacutainer tubes. After inversion (10 times), we centrifuged the vacutainer tubes at 4 °C for 10 minutes at 1500 g without brakes. The plasma is then transferred into a 15 mL centrifuge tube and centrifuged for a second time for 10 minutes at 1500 g. Finally, the plasma was transferred into cryovials and stored at −80 °C until shipment. The cancer types included are colorectal cancer (CRC), lung adenocarcinoma (LUAD), and prostate cancer (PRAD).

#### RNA isolation and spike-in controls

Total RNA was isolated from platelet-free (PFP) and platelet-rich plasma (PRP) using the miRNeasy Serum/Plasma Kit (Qiagen, Hilden, Germany, 217184). We used 200 μl of plasma as input. For the cancer patient experiment, 2 μl of 1x RC PFP spikes were added to the plasma during isolation. The elution volume was 14 μl, and we added 2μl of 1x LP PFP spikes (Thermo Fisher). Detailed descriptions of the spike-in controls can be found in the exRNAQC study (Consortium et al., 2021). From this total volume, we used 5 μl for the library preparation. For the healthy donor experiment, the eluate of multiple parallel extractions was pooled according to the original biofluid type (PRP or PFP) and split into six aliquots of 5 μl this to minimize extraction bias. We did not include a gDNA removal step after RNA isolation. The input is volume-based since the RNA concentrations of PFP and PRP are below the limit of quantification.

#### YRNA LNA design

The YNRA4 fragment (32 nucleotides) was tiled with 16 bp long complementary nucleotides resulting in 17 possible designs. We mapped the full set of antisense oligonucleotides to the human transcriptome (Ensembl v84) and miRBase. Oligonucleotides with no off-targets when 3 mismatches are allowed were retained. Of the retained LNAs, we chose the oligonucleotide with the highest melting temperature (Tm). The resulting fully modified LNA (ACCCACTACCATCGGA, targeting TCCGATGGTAGTGGGT) has a Tm of 89.9 °C. In addition to the fully LNA-modified oligo, for the same sequence we ordered 2’-O-methyl and 2’-methoxy-ethoxy modified nucleotides and half modified (alternating modified – non-modified nucleotides) oligos at Integrated DNA Technologies. Sequences are available in Supplemental Table 1.

#### TruSeq small RNA library prep

We used the TruSeq small RNA library prep sequencing kit (Illumina, San Diego, CA, USA) for library preparation according to manufacturing instructions except for the changes listed below. After adaptor ligation and before the reverse transcription step, 2 μl LNA with a concentration of 0.25 μM (LNA1x) or 2.5 μM (LNA10x) was added to 14 μl of the adaptor-ligated RNA. In the experiments with the cancer patient samples and alternative modifications, only the 0.25 μM (LNA1x) concentration was analyzed as we showed that the 10-fold higher concentration had no added value. As a negative control for LNA blocking (LNA0x), 2 μl of water was added to 14 μl of RNA. Next, we used 6 μl of each sample to start the reverse transcription and continue the library prep. Since the input amounts are low, the number of PCR cycles was set at 16 (the manufacturer recommends 11) during the final PCR step.

#### Pippin prep and sequencing

We performed a size selection for 125–163 bp on all libraries using 3% agarose dye-free marker H cassettes on a Pippin Prep (Sage Science, Beverly, MA, USA). Next, the libraries were purified by precipitation using ethanol and resuspended with 10 mM Tris-HCl buffer (pH 8.0) with Tween 20. After dilution, the libraries were quantified using the KAPA Library Quantification Kit (Roche Diagnostics, Diegem, Belgium, KK4854). Healthy donor samples were sequenced using a NextSeq 500 using the NextSeq 500 High Output Kit v2.5 (75 cycles) (Illumina, San Diego, CA, USA). We loaded the library at a concentration of 2.0 pM with 10% PhiX and obtained a total of 268 M reads. We loaded the cancer patient samples on one lane of a NovaSeq 6000 (Illumina, San Diego, CA, USA) instrument at a concentration of 300 pM with 10% PhiX using the NovaSeq 6000 SP Reagent Kit v1.5 (100 cycles) (Illumina, San Diego, CA, USA) (paired-end, 2x 50 cycles, only the first read was used for subsequent analysis), resulting in 267M reads. For the chemical modification comparison experiment, we used one lane of a NovaSeq 6000 SP Reagent Kit v1.5 (100 cycles) (Illumina, San Diego, CA, USA, 20028401) (Illumina, San Diego, CA, USA) (1×100 bp), loading 300 pM with 10% PhiX, resulting in a total of 548M reads.

#### Quantification analysis

We used a dedicated in-house small RNA-seq pipeline for the quantification of small RNAs. This pipeline starts with adaptor trimming using Cutadapt (v1.8.1) (Martin, 2011), which discards reads shorter than 15□nt, and those in which no adaptor was found. The reads with a low quality are discarded by using the FASTX-Toolkit (v0.0.14) (“FASTX-Toolkit,” n.d.) set at a minimum quality score of 20 in at least 80% of nucleotides. Next, we counted and filtered out reads belonging to our spike-in controls (both RC as LP). The spike reads are subtracted from the FASTA files, and reads are counted. For this comparison, the spike-in controls were not used for correction since the library preparation methods (adding LNA or not) differ. The spike-ins are, however, needed to correct for input concentration variation when all other parameters are equal, as the pooling is performed based on volume. Subsequently, we mapped the reads with Bowtie (v1.1.2) (Langmead et al., 2009), allowing one mismatch. At the end of the pipeline, the mapped reads are annotated by matching the genomic coordinates of each read with genomic locations of miRNAs (obtained from miRBase, v20) and other small RNAs (obtained from UCSC GRCh37/hg19 and Ensembl v84). We submitted the original FASTQ-files and the count tables in EGA (EGAS00001006023). The samples are downsampled to the sequencing depth of the sample with the least number of reads per experiment, or respectively 13M reads (concentration experiment), 6.5M reads (modification experiment), and 7M reads (cancer experiment).

#### Computational analysis

We used R (v3.6.0) (R Core Team, 2021) for further data processing, using the following packages: tidyverse (v1.2.1) (Wickham et al., 2019), biomaRt (v2.40.4) (Durinck et al., 2009, 2005), broom (v0.5.2) (Robinson, 2014). For differential expression analysis limma-voom (v3.40.6) (Ritchie et al., 2015) was used on a filtered matrix with at least 10 reads per million (RPM) per miRNA over all samples.

### Mitochondrial ribosomal RNA blocking in cell lysates

#### Cell culture and RNA extraction

We used HEK293T cells that were grown in RPMI 1640 medium with GlutaMAX supplement (Thermo Fisher, Waltham, MA, USA) supplemented with 10% fetal calf serum (Merck, Germany) and were lysed with SingleShot lysis buffer (Bio-Rad, United States of America).

#### MtRNA LNA design

From previous experiments, we identified three transcripts without poly(A) tail that are abundant (0.1-2% of all counts) in 3’ end sequencing data of HEK293T cells: MT-RNR1, MT-RNR2, and RNA45S. We visually inspected the RNA sequencing data using IGV_2.7.2 (Robinson et al., 2011) and confirmed the presence of an adenosine-rich region flanking the abundant fragments observed in the sequencing library. For MT-RNR2, two different fragments were associated with an internal poly(A) stretch, contributing to the high number of gene counts. We investigated a design region of about 50 bases overlapping the abundant fragments and used Bowtie (v1.2.3) (Langmead et al., 2009) to map several 16-base-long putative LNA sequences. We retained the oligos with the lowest number of off-target hits. We then checked their binding capacities and biochemical characteristics. Sequences are available in Supplemental Table 1.

#### LNA treatment

We combined four different LNA mixes (MT-RNR2_1, MT-RNR2_2, MT-RNR1, and RNA24S) to have a final solution containing each LNA at 25 μM (100x). We mixed 2 μl of LNA to 3 μl of RNA sample. From this solution, we used 2.5 μl as input for the library preparation.

#### Library preparation

For the library preparation, we used the QuantSeq 3’ mRNA-Seq Library Prep Kit FWD for Illumina (Lexogen, Austria). We performed the ‘low input’ version of the protocol.

#### Sequencing

We sequenced the libraries using the NovaSeq 6000 SP Reagent Kit v1.5 (100 cycles) (Illumina, San Diego, CA, USA) on a NovaSeq 6000 (Illumina, San Diego, CA, USA) instrument at a concentration of 300 pM with 10% PhiX.

#### Quantification analysis

We used BBMap (v38.26) to trim off the poly(A) tails and adapter sequences and to perform quality trimming. Next, all FASTQ files were subsampled to 2,000,000 reads with Seqtk (v1.3) and mapped to the hg38 genome using STAR (v2.6.0). We used SAMtools (v1.9) to count the reads mapping to the LNA-targeted genomic regions. We used htseq-count (v0.11.0) (Anders et al., 2015) to generate the overall counts. Before initial trimming, before quality trimming, and after quality trimming, we used FastQC (v0.11.9) to investigate the quality of the reads.

#### Computational analysis

We used R (v4.1.0) (R Core Team, 2021) and tidyverse (v1.3.1) (Wickham et al., 2019) and biomaRt (v2.48.3) (Durinck et al., 2009, 2005) to analyze and visualize the computationally generated data.

### Mitochondrial ribosomal RNA blocking in direct-cDNA long-read sequencing

#### Cell culture and harvesting

We cultured HEK293T cells in RPMI medium supplemented with 10% fetal calf serum to 80% confluence in a T75. The cells were washed with 2 ml versene and incubated with 2 ml of trypsin for 3 minutes at 37 °C. We neutralized the mixture with 8 ml fresh medium. We centrifuged for 5 minutes at 2000 rcf at 4 °C and removed the supernatants. We resuspended the cells in 1 ml of QIAzol and flash-froze the mixture in liquid nitrogen.

#### RNA extraction and quality control

We extracted RNA using the RNeasy Micro Kit (Qiagen, Hilden, Germany, 217184) according to the manufacturer’s protocol. We checked the quality of the RNA (RQN = 10) using a Fragment Analyzer (Agilent, United States of America).

#### LNA treatment

We combined four different LNA mixes (MT-RNR2_1, MT-RNR2_2, MT-RNR1 and RNA24S) to have a final solution containing each LNA at 25 μM. We then made a 10-fold dilution series to obtain three different LNA solutions: LNA1x (0.25 μM), LNA10x (2.5 μM) and LNA100x (25 μM). For each library preparation, 2 μg of total RNA was mixed with 2 μl of the corresponding LNA dilution. 1 μl of RNase-free water was added to 2 μg of total RNA as a non-treated sample (LNA0x). The samples are place on ice for 5 minutes.

#### Library preparation

We prepared direct-cDNA libraries using the SQK-DCS109 Kit (Oxford Nanopore Technologies, United Kingdom). The exact protocol was followed except for the following changes: the RNA-bead binding steps were performed for 5 minutes on a Hula Mixer and 5 minutes on the bench at room temperature; the RNA elution steps were performed for 5 minutes at 37 °C and 5 minutes on a Hula Mixer at room temperature; and 300 μl of 80% ethanol was used for the beads wash steps.

#### Oxford Nanopore sequencing

We sequenced each library using two Flongle Flow Cells (Oxford Nanopore Technologies, United Kingdom) with a MinION Sequencer (Oxford Nanopore Technologies, United Kingdom). Sequencing was either stopped after 24 hours or when no more pores were available.

#### Quantification analysis

We basecalled the raw fast5 files using Guppy (v3.5.2) (Wick et al., 2019) on a GPU. We grouped reads per sample and used Pychopper (v2.3.1) (“nanoporetech/pychopper: A tool to identify, orient, trim and rescue full length cDNA reads,” n.d.) to identify full-length transcripts containing both primer sequences. We mapped the reads with Minimap2 (v2.11) (Li, 2018) and extracted reads mapping to the target fragment location using SAMtools (v1.11) (Danecek et al., 2021). We then used NanoComp (v1.12.0) (De Coster et al., 2018) to check the read length and quality of each sample.

#### Computational analysis

We used R (v4.1.0) (R Core Team, 2021) and tidyverse (v1.3.1) (Wickham et al., 2019) and biomaRt (v2.48.3) (Durinck et al., 2009, 2005) to analyze and visualize the computationally generated data.

### MALAT1 blocking in single-cell 3’ end sequencing for *peripheral blood mononuclear cells* (PBMCs)

#### PBMCs preparation

We collected whole blood in EDTA tubes. The blood was transferred to Leucosep filtered tubes (Greiner Bio-One) containing 15 ml of Ficoll Paque Plus (Cytiva, Washington, D.C., USA, 17144002) and diluted (1:2) with the same volume of 1X DPBS (Thermo Fisher, Waltham, MA, USA, 14190144). We centrifuged the samples at room temperature for 18 minutes at 800 rcf and extracted the PBMCs from the resulting buffy coat. The extracted PBMCs were centrifuged and washed twice with 1X DPBS (Thermo Fisher, Waltham, MA, USA, 14190144). We took a sample for counting, and assessed the cell viability and concentration with a Neubauer chamber, counting at least two different squares. PBMCs were then resuspended in freezing mix (complete medium (RPMI + 1% pen/strep + 10% FCS) + 10% DMSO) in cryovials with no more than 10 million cells. The vials were stored first at −80 °C inside a freezing container for 24 h and then at −150 °C. We thawed the vials just before live-death sorting.

#### MALAT1 LNA design

After visually inspecting 3’ end sequencing data from PBMCs using IGV_2.7.2 (Robinson et al., 2011), the optimal design space was identified (Supplemental Figure 8). We identified two internal poly(A) sequences contributing to the high number of counts. Next, we designed and characterized the best LNA sequences following similar steps as before (see ‘mtRNA LNA design’, but with a length of 18 nucleotides). The sequences are available in Supplemental Table 1.

#### LNA treatment

We diluted the LNAs at 125 μM of which 2 μl was used. This concentration is higher than the YRNA experiment, as we expect the total RNA concentration to be higher for this experiment. For the pre-RT blocking, 2 μl of the oligonucleotide mix was added to the master mix (including the RT reagent, template switching oligo, reducing reagent B, and RT enzyme C). The master mix is then combined with the cell suspension to a total volume of 80 μl. For the pre-cDNA amplification blocking, we added 2 μl of the oligonucleotide mix to the cDNA amplification mix (including Amp Mix and cDNA primers).

#### Library preparation

Sorted single-cell suspensions were resuspended in PBS+0.04% BSA at an estimated final concentration of 1000 cells/μl and loaded on a Chromium GemCode Single Cell Instrument (10x Genomics, Pleasonton, CA, USA, 1000204), Chip G (10x Genomics, Pleasonton, CA, USA, #2000177) to generate single-cell gel beads-in-emulsion (GEM). We prepared the scRNA-seq libraries using the GemCode Single Cell 3’ Gel Bead and Library kit, version NextGEM 3.1 (10x Genomics, Pleasonton, CA, USA, PN-1000121) according to the manufacturer’s instructions.

#### Sequencing

The Chromium libraries were equimolarly pooled and loaded on a NovaSeq 6000 (Illumina, San Diego, CA, USA) instrument in standard mode with a final loading concentration of 340 pM and 2% PhiX. We obtained a total of 952M reads with q30 of 91.32% with an SP100 cycles (Illumina, San Diego, CA, USA, 20028401) kit. The number of (pre-filtered) cells per experiment were highly comparable, 13841 cells for the noLNA sample, 13279 cells pre-RT, and 13893 cells post-RT. The FASTQ files were subsampled based on the number of cells to obtain a comparable number of reads/cell over all samples.

#### Quantification analysis

Demultiplexing of the bcl files was performed with cellranger mkfastq (v6.0.1), after which gene counts per cell were obtained with cellranger count (v6.0.1).

#### Computational analysis

The count matrixes were loaded into R (v4.1.0) (R Core Team, 2021) and further processed, including the integration and annotation, with Seurat (v4.0.3) (Hao et al., 2021). We did not filter the cells. We analyzed and visualized the data using tidyverse (v1.2.1) (Wickham et al., 2019).

### LNA blocking simulation in whole blood 3’-end sequencing

#### Data download

We downloaded one of the whole blood 3’-end RNA sequencing (QuantSeq) samples generated by Uellendahl-Werth et al. (Uellendahl-Werth et al., 2020) (SRR11028518). This sample had a sequencing depth of 18,043,131 reads.

#### Quantification analysis

We used BBMap (v38.26) to trim off the poly(A) tails and adapter sequences and to perform quality trimming. Next, we mapped the trimmed reads to the hg38 genome using STAR (v2.6.0). We used htseq-count (v0.11.0) (Anders et al., 2015) to quantify the uniquely mapped reads. We used FastQC (v0.11.9) to investigate the quality of the reads before quality trimming and after quality trimming.

#### Depletion simulations

All simulations were run using R (4.1.0). First, we generated the sampling distribution by first removing the ENSG00000244734 (HBB) reads and calculating the fraction of reads appointed to each gene relative to the total amount of reads. We used this distribution to guide the subsampling. We then subsampled the count tables for a varying total number of counts (0.5M, 1M, 2M, 4M, 8M), initial HBB abundance (0-90%, by 10% increments), and percentage of depletion (0-100%, by 2% increments). Last, we calculated the number of genes with 10 counts or larger. Finally, we analyzed and visualized (v1.2.1) (Wickham et al., 2019).

## RESOURCE LIST

- 1X DPBS (Thermo Fisher, Waltham, MA, USA, 14190144).
- RPMI 1640 medium with GlutaMAX supplement (Thermo Fisher, Waltham, MA, USA, 61870010
- 10% Fetal calf serum (Merck, Germany, F0804-500ML)
- Chromium GemCode Single Cell Instrument (10x Genomics, Pleasonton, CA, USA, 1000204)
- Direct cDNA sequencing kit (Oxford Nanopore Technologies, UK, SQK-DCS109)
- Eppendorf Protein LoBind microcentrifuge tubes (Eppendorf, Hamburg, Germany, Z666548)
- Ficoll Paque Plus (Cytiva, Washington, D.C., USA, 17144002
- Flongle Flow cell (Oxford Nanopore Technologies, UK, FLO-FLG001)
- Fragment Analyzer RNA Kit (Agilent, USA, DNF-471-0500)
- GemCode Chip G (10x Genomics, Pleasonton, CA, USA, 2000177)
- GemCode Single Cell 3’ Gel Bead and Library kit, version NextGEM 3.1 (10x Genomics, Pleasonton, CA, USA, PN-1000121)
- KAPA Library Quantification Kit (Roche Diagnostics, Diegem, Belgium, KK4854)
- MinION sequencer (Oxford Nanopore Technologies, UK, MIN-101B)
- miRNeasy Serum/Plasma Kit (Qiagen, Hilden, Germany, 217184).
- NextSeq 500 High Output Kit v2.5 (75 cycles) (Illumina, San Diego, CA, USA, 20024906)
- NextSeq 500 Sequencing System (Illumina, San Diego, CA, USA, SY-415-1001)
- NovaSeq 6000 Sequencing System (Illumina, San Diego, CA, USA, 20012850)
- NovaSeq 6000 SP Reagent Kit v1.5 (100 cycles) (Illumina, San Diego, CA, USA, 20028401)
- Pippin Prep (Sage Science, Beverly, MA, USA, PIP0001).
- QuantSeq 3’ mRNA-Seq Library Prep Kit FWD for Illumina (Lexogen, Austria, 139.96)
- RNeasy Micro Kit (Qiagen, Hilden, Germany, 217184)
- SingleShot lysis buffer (Bio-Rad, United States of America, 1725080)
- TruSeq small RNA library prep sequencing kit (Illumina, San Diego, CA, USA, RS-200)
- Vacutainer Hemogard Closure Plastic K2-Edta Tube, 10 ml, (BD, Franklin Lakes, NJ, USA, 367525)
- Vacutainer Push blood collection set (BD, Franklin Lakes, NJ, USA, 368657)
- HEK293T (ATCC, Manassas, VA, USA)

## RESULTS

To prevent the incorporation of unwanted (fragments of) transcripts in RNA sequencing libraries, we reasoned that LNA-modified oligonucleotides would block reverse transcription and PCR amplification when bound downstream of the priming site because of their extremely high affinity to RNA and cDNA. The approach we took to design blocking LNA oligonucleotides depends on the characteristics of the unwanted RNA sequence and the library prep procedure. We selected four library prep procedures and defined high abundant and mostly unwanted target RNA sequences for LNA oligonucleotide design (Figure 1A). These targets include YRNA in small RNA sequencing libraries from human blood plasma, mitochondrial rRNA in 3’ end sequencing libraries and long read sequencing libraries of HEK293T cells, and MALAT1 in single-cell 3’ end sequencing libraries of PMBCs. To block RT and PCR of short fragments like YRNA in small RNA-seq libraries, we designed an 18 nucleotide LNA to be complementary to the 3’ end of the 30 nucleotide YRNA fragment (Figure 1B). For longer fragments, like mitochondrial rRNA and MALAT1, the LNA oligonucleotide was designed to bind directly downstream of the poly(A) RT-priming site (Figure 1B-F). As the LNA oligonucleotides are added directly to the RT reaction (see details in Material and Methods for each of the protocols), no additional steps are required in the RNA library prep protocol. Since the LNA remains present during the PCR steps, it will also inhibit the remaining fragments during PCR amplification (Figure 1G).

**Figure 1.**
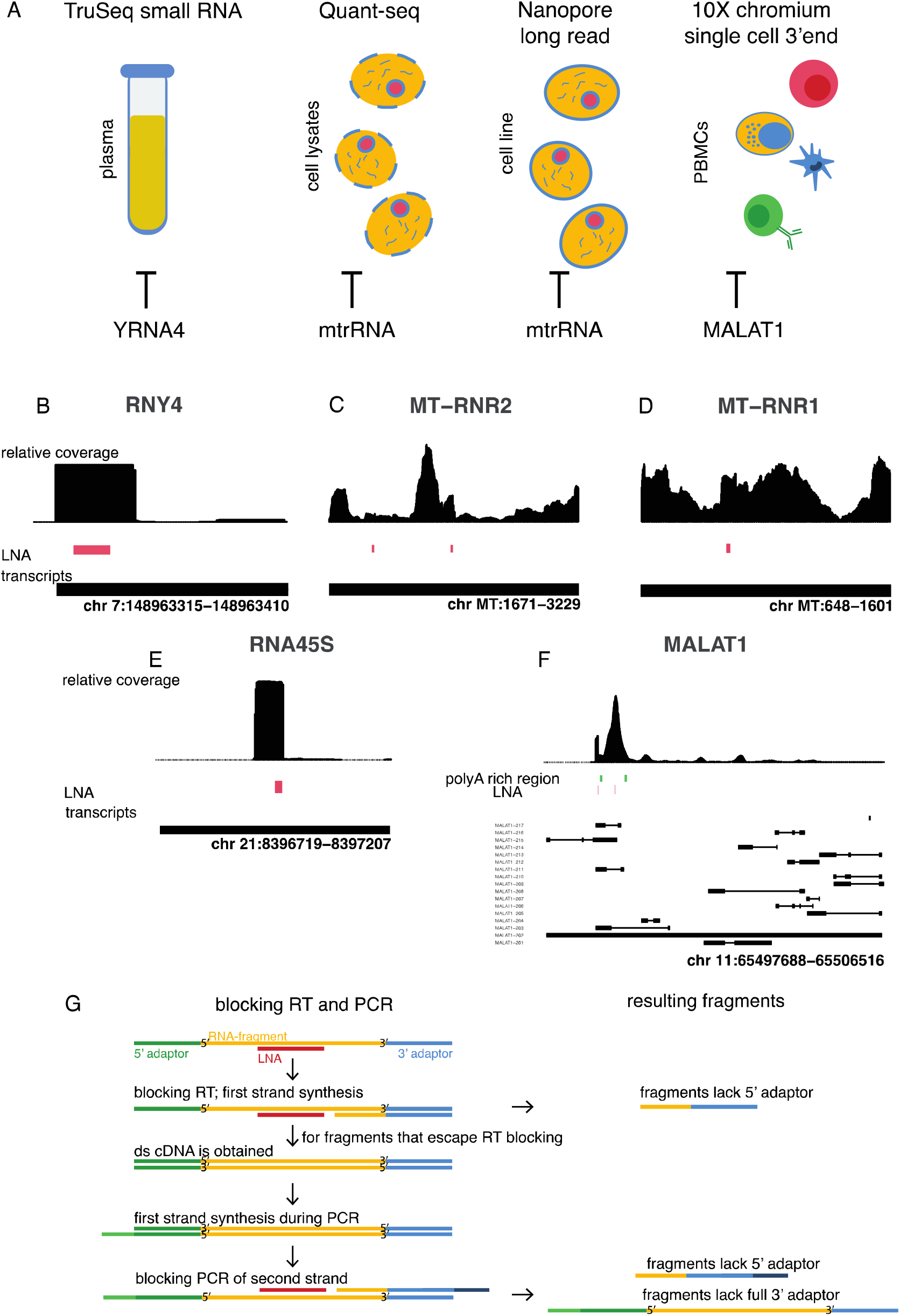
Overview of targeted transcripts and investigated library preparation methods. A. Visual representation of the evaluated RNA sequencing library preparation methods with the associated blocked transcripts. B-F. Coverage plots of standard library prep RNA-sequencing data for the targeted region. The height of the bars represents the relative number of reads mapping to that position in the transcript. The LNA oligonucleotide and its binding location is shown in red/green under the coverage plot. The targeted transcripts and their chromosomal location are indicated at the bottom of each plot. G. Schematic overview of blocking during RT and PCR amplification.

### YRNA blocking in plasma samples for TruSeq small RNA sequencing

#### Efficient blocking of RNY4 in PRP and PFP

We first focused on blocking RT and amplification of RNY4 fragments in human blood plasma small RNA-seq libraries. We tested the blocking efficiency on platelet-rich plasma (PRP) and platelet-free plasma (PFP) from healthy donors, with PRP having the highest fraction of RNY4 fragments (PFP: x%, PRP: y%). We then spiked two different concentrations of a blocking LNA oligonucleotide (0.25 μM and 2.5 μM, referred to as LNA1x and LNA10x, respectively) in the RT reaction of the TruSeq small RNA library prep, and compared the results to the standard workflow. We only observed 0.09% RNY4 in PFP and up to 0.16% RNY4 in PRP (Figure 2A) when adding LNA1x to the RT reaction, or respectively a 477- and 468-fold reduction compared to the standard protocol. Increasing the LNA concentration 10-fold (LNA10x) provided no benefit compared to LNA1x, with a 228-fold and 262-fold reduction of RNY4. The strong reduction in RNY4 fragments was accompanied by a strong increase in fraction of microRNA reads, from 49.55% to 79.67% for PFP and from 17.24% to 74.61% for PRP. Since the LNA1x condition resulted in a sufficient reduction in YRNA and an increase in microRNA read fraction, we decided to use the 1x concentration (0.25 μM) for the subsequent experiments unless specified otherwise.

**Figure 2.**
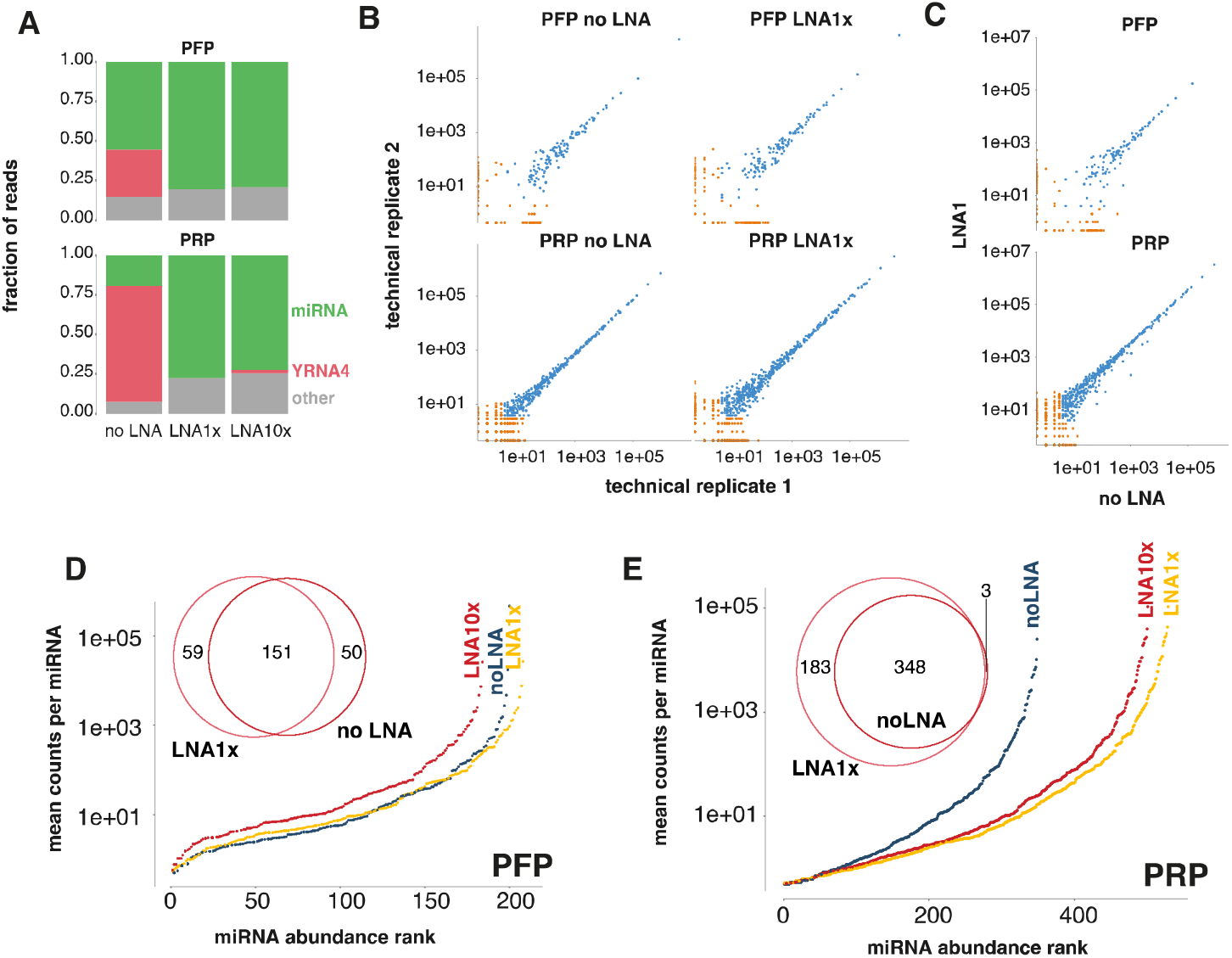
Blocking of RNY4 transcripts in platelet-rich and platelet-free plasma using different blocking LNA concentrations. A. Read percentages for the YRNA and miRNA biotype without the LNA (noLNA) blocking and by adding the LNA in two different concentrations (LNA1x, LNA10x) to both platelet rich (PRP) and platelet free (PFP) plasma. B. Correlation plots between technical replicates for the blocking strategies and samples. Each dot represents a miRNA. The dots are colored orange when the miRNA had less than 5 counts in either of the replicates. C. Correlation plots of the counts between treated (LNA1x) and untreated (no LNA) samples. Only technical replicate 1 is shown here due to the high technical reproducibility. D & E. The mean counts per million (CPM) over the technical replicates for all detected (CPM > 0.5) miRNAs and overlap between the detected miRNAs in the LNA blocking (LNA1x) and no blocking condition (noLNA) for both PFP and PRP.

#### RNY4 blocking increases microRNA coverage and preserves fold changes

We then evaluated the reproducibility of our RNY4 blocking protocol by comparing miRNA abundance between technical library preparation replicates. Reproducibility upon RNY4 blocking was similar to that of the standard workflow, as evidenced by similar Pearson (0.999-1) and Spearman (0.70-0.78 for PFP and 0.81-0.82 for PRP) correlation coefficients (Figure 2B). To investigate if the abundance of miRNAs is affected by RNY4 blocking (due to off-target effects), we compared miRNA abundance (reads per million) between the RNY4 blocking procedure and the standard workflow. Only one miRNA showed a high standardized residual (residual divided by standard deviation) (>2) in all samples and replicates. This miRNA (miR-106b-3p) showed a consistently lower abundance in the LNA1x libraries compared to the control. Of note, we did not observe any sequence similarity between the RNY4 fragment and miR-106-3p, suggesting that non-specific binding of the LNA is unlikely. In general, miRNA expression correlations between the standard protocol and LNA1x spike protocol (Figure 2C) were comparable to these of technical replicates (Pearson correlation = 1.00, Spearman correlation = 0.67-0.72 for PFP and 0.81 for PRP). In PFP, the impact of RNY4 blocking on the number of detected miRNAs was limited, with only nine additional miRNAs detected upon subsampling for library size correction (Figure 2D). However, the coverage for the detected miRNAs increased (Supplemental Figure 1B). As expected, the uniquely detected miRNAs were low abundant (Supplemental Figure 1A&1B). In PRP, we detected 183 additional miRNAs in the LNA1x spike protocol. All except three miRNAs detected with the standard protocol were also detected with the LNA1x spike protocol (Figure 2E, Supplemental Figure 1C). Not only does RNY4 blocking increase the number of detected miRNAs, it also results in increased miRNA coverage (a 3-fold median RPM increase) (Supplemental Figure 1D). Taken together, RNY4 blocking in blood plasma small RNA sequencing libraries improves miRNA library complexity and coverage.

We then assessed the impact of RNY4 blocking on differential miRNA abundance between samples. To address this, we examined the miRNA fold changes between PFP samples from patients with diverse tumor types (colorectal cancer or CRC (n=4), prostate adenocarcinoma or PRAD (n=4) and lung adenocarcinoma or LUAD (n=4)) that were processed with the standard and LNA1x spike protocol (Supplemental Figure 2). Differences in miRNA abundance between cancer types were highly concordant between both methods (Figure 3A). To further assess the impact of RNY4 depletion on the robustness of differential expression analysis at various sequencing depths, we repeatedly subsampled reads to various sequencing depths and determined the concordance of differential miRNAs between the different subsamples. At high sequencing depth (7M reads), both methods result in an equal concordance between the differentially expressed miRNAs amongst subsamples. When subsampling reads to a lower sequencing depth, however, the concordance between detected differential miRNAs (i.e., the number of shared differential miRNAs between subsamples) was higher in the case of RNY4 blocking (Figure 3B). At 0.7M reads, the average concordance of four random subsamples was 64.9% for RNY4 blocking and 60.0% without RNY4 blocking. When using 7M reads, the concordance was 100% for both methods. This observation suggests that, at shallow sequencing depth, RNY4 blocking increases the robustness of differential miRNA analysis.

**Figure 3.**
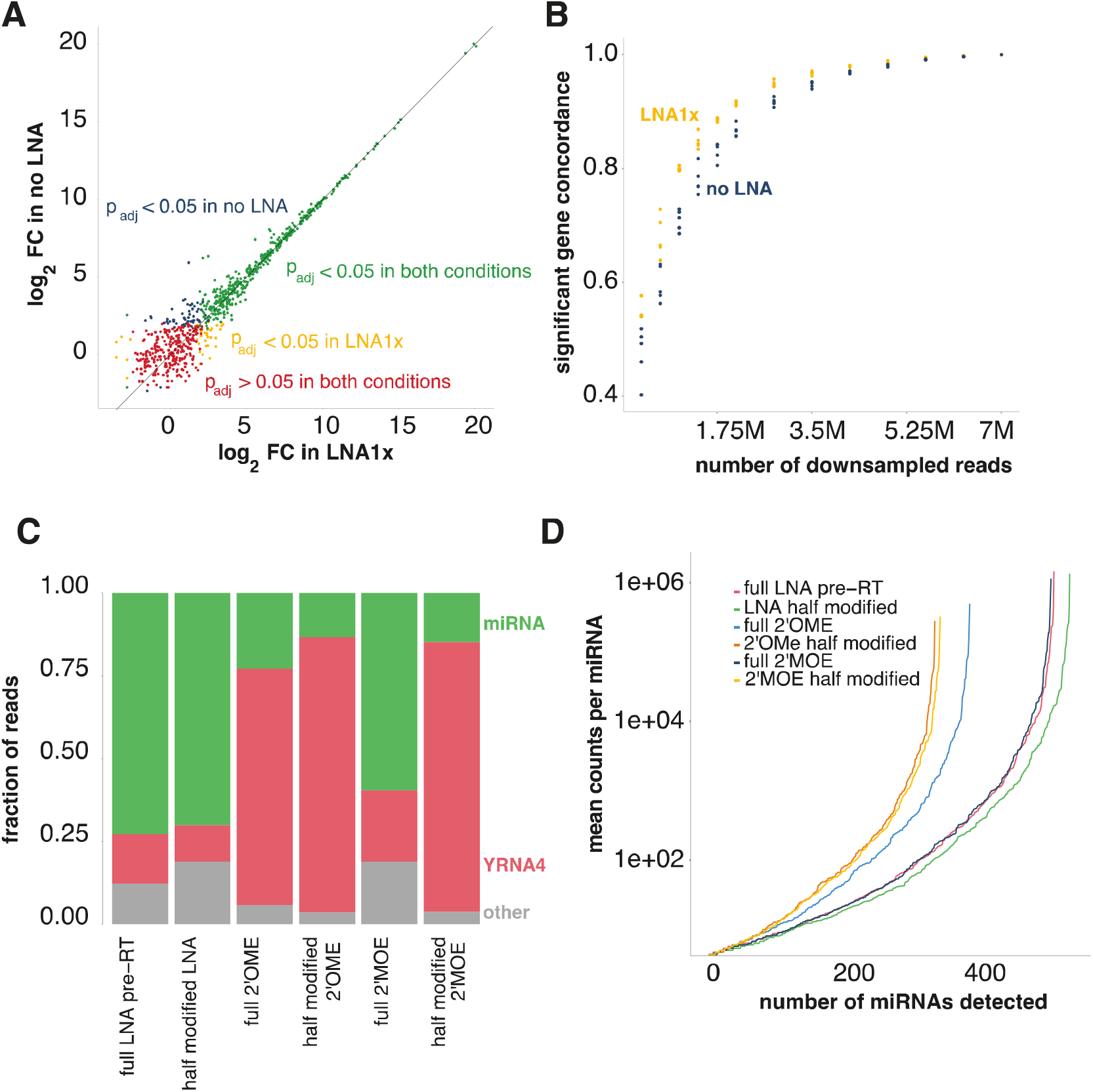
Depletion results for alternative blocking oligonucleotide modifications. A. Correlation plot between the log_2_ fold changes of noLNA and LNA1x samples between PRAD and CRC samples. Each dot is a miRNA and is colored by its p-value in the treated and untreated samples. B. Significant gene concordance (i.e., the number of overlapping DE miRNAs between subsamples, as a measure for robustness) for four subsamples per sample type (noLNA and LNA1x) for different subsample sizes. C. Read percentages for the YRNA and miRNA biotype for multiple modifications. D. The number of discovered miRNAs and their coverage for the various modifications.

#### LNA is most efficient modification to block library incorporation

As fully modified LNA oligonucleotides are relatively expensive, we evaluated the RNY4 blocking potency of cheaper base modifications known to improve oligonucleotide binding affinity such as 2’-O-methyl (2’OME) and 2’-methoxy-ethoxy (2’MOE) in PRP. We observed that an LNA-modified RNY4 oligonucleotide is equally efficient (median reduction of 6.45-fold) compared to a 2’MOE modified oligonucleotide (median reduction of 6.11-fold) (p=0.14). The 2’OME modification, however, is less efficient and resulted in reduction of just 1.22-fold (p=0.005) compared to the LNA modification. In addition, we investigated the potency of partially modified (i.e., every other nucleotide) oligonucleotides for both LNA, 2’OME, and 2’MOE. We observed that the partially modified LNA RNY4 oligonucleotide was as potent as a fully modified LNA RNY4 oligonucleotide (RNY4 fold change reduction of 6.90, p=0.383) and still outperforms fully modified 2’OME RNY4 oligonucleotides (p=0.0005) (Figure 3C). The partially 2’OME and 2’MOE modified oligonucleotides performed the worst (Figure 3C).

### rRNA blocking in 3’ end sequencing data

During reverse transcription, oligo(dT) primers can bind internal poly(A) sequences of mitochondrial and nuclear ribosomal RNA species, which eventually get incorporated in the RNA sequencing library (up to 2% of all reads, as found in previous sequencing data (Figure 1C-E)). Although the abundance of MT-rRNA in this data is not necessarily problematic, it does provide a good test case for eliminating multiple transcript fragments in a poly(A)-primed library prep procedure. Therefore, we designed fully modified LNA oligonucleotides to inhibit the reverse transcription of three mitochondrial rRNA fragments (MT-RNR1 and two fragments from MT-RNR2) and one fragment from nuclear rRNA RNA45S (Figure 1). We added all four oligonucleotides to the RT reaction of a 3’ end library preparation on eight cell lysates and compared the data to that of a standard 3’ end library preparation workflow. Adding LNA oligonucleotides resulted in an average reduction of the counts per million of 16.2x, 19.2x, 8.6x, and 3.2x for RNA45S, MT-RNR1, MT-RNR2 fragment 1, and MT-RNR2 fragment 2, respectively (Figure 4A). To evaluate the reproducibility of the method, we compared the abundance of all detected genes between biological replicates for both the standard and the LNA spike protocols. The Pearson (0.982-0.997) and Spearman (0.852-0.879) correlation coefficients were high for every comparison (Supplemental Figure 3), and there was no significant difference in reproducibility between the standard and blocking protocol. We evaluated the number of detected genes for different sequencing depths to investigate whether blocking RNA45S and the MT-RNR1/2 fragments is beneficial for gene detection (Figure 4B). For shallow sequencing depth (1-2 million reads), the number of detected genes was higher in the blocking protocol compared to the standard protocol. Finally, we investigated the potential off-target effects of the blocking oligonucleotides by comparing gene expression values between the control and blocking protocol. Out of 12,077 detected genes, we identified two genes that showed divergent gene expression values for all biological replicates: MT-ATP8 and H4C3 (Figure 4C). We did not observe significant sequence complementarity between the LNA oligonucleotides and these presumed off-targets. In conclusion, LNA oligonucleotides can efficiently and specifically block the incorporation of a variety of transcript fragments in 3’ end RNA-seq libraries.

**Figure 4.**
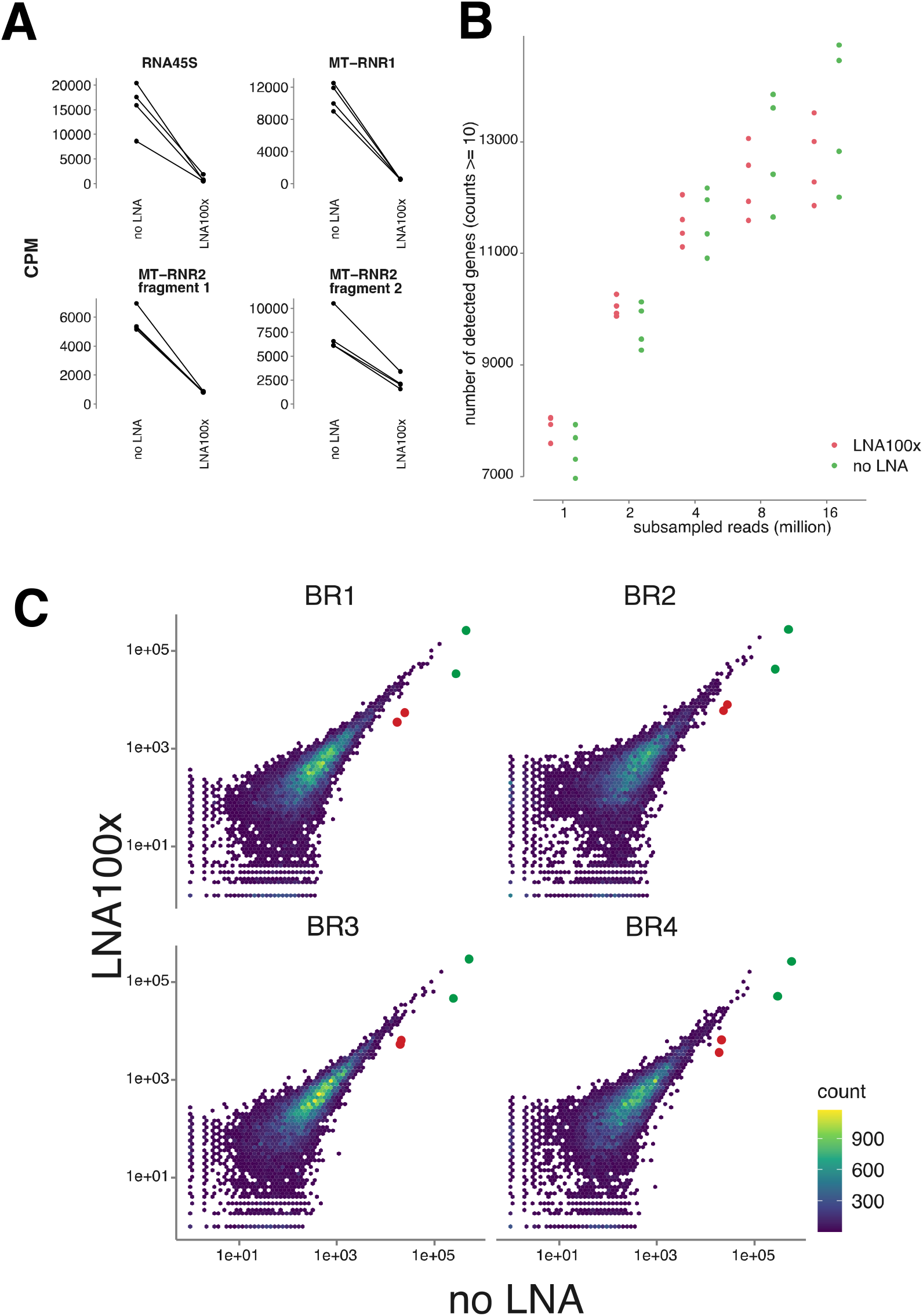
LNA oligonucleotide blocking of mtrRNA fragments (mitochondrially encoded 16S rRNA (two fragments), 45S pre-ribosomal RNA and mitochondrially encoded 12S rRNA) in QuantSeq 3’ end sequencing. A. Counts per million (CPM) of the targeted transcripts for noLNA and LNA100x samples after QuantSeq 3’ end sequencing. Lines are drawn between samples originating from the same lysate. B. Average number of detected genes for subsampled noLNA and LNA100x samples. A gene is ‘detected’ if it has at least 10 counts. Each point is a biological replicate and is colored by treatment. C. Scatter plot between noLNA and LNA100x samples for each biological replicate are highly correlated (Spearman = 0.853-0.905). Each dot corresponds to a gene. The green dots are MT-RNR1 and MT-RNR2, while the red dots indicate the genes likely affected by off-target binding: H4C3 and MT-ATP8.

### rRNA blocking for long-read polyA+ transcript sequencing

Additionally, we explored whether the previously described rRNA (RNA45S, MT-RNR1, MT-RNR2) blocking strategy can also be applied to Oxford Nanopore Technologies (ONT) sequencing of poly(A)-primed cDNA libraries. More specifically, we performed direct-cDNA sequencing to investigate the blocking effect on just the reverse transcription step. We added three different concentrations (0.25 μM, 2.5 μM, and 25 μM, referred to as LNA1x, LNA10x, and LNA100x) of the rRNA LNA oligonucleotides (as used in the 3’ end library preparation) to the reverse transcription reaction of four different samples. For all targeted fragments, we observed a substantial decrease in counts per million with increasing concentration of LNA oligonucleotides, except for 45S pre-ribosomal RNA in the LNA100x condition (Figure 5). Unexpectedly, we also observed a mild but consistent decrease in overall read length distribution with increasing concentration of LNA oligonucleotides (Supplemental Figure 4). The quality scores of the reads did not vary (Supplemental Figure 5). These results show the potential of LNA oligonucleotides to prevent reverse transcription (and thus sequencing) of specific RNA molecules in ONT long-read sequencing experiments.

**Figure 5.**
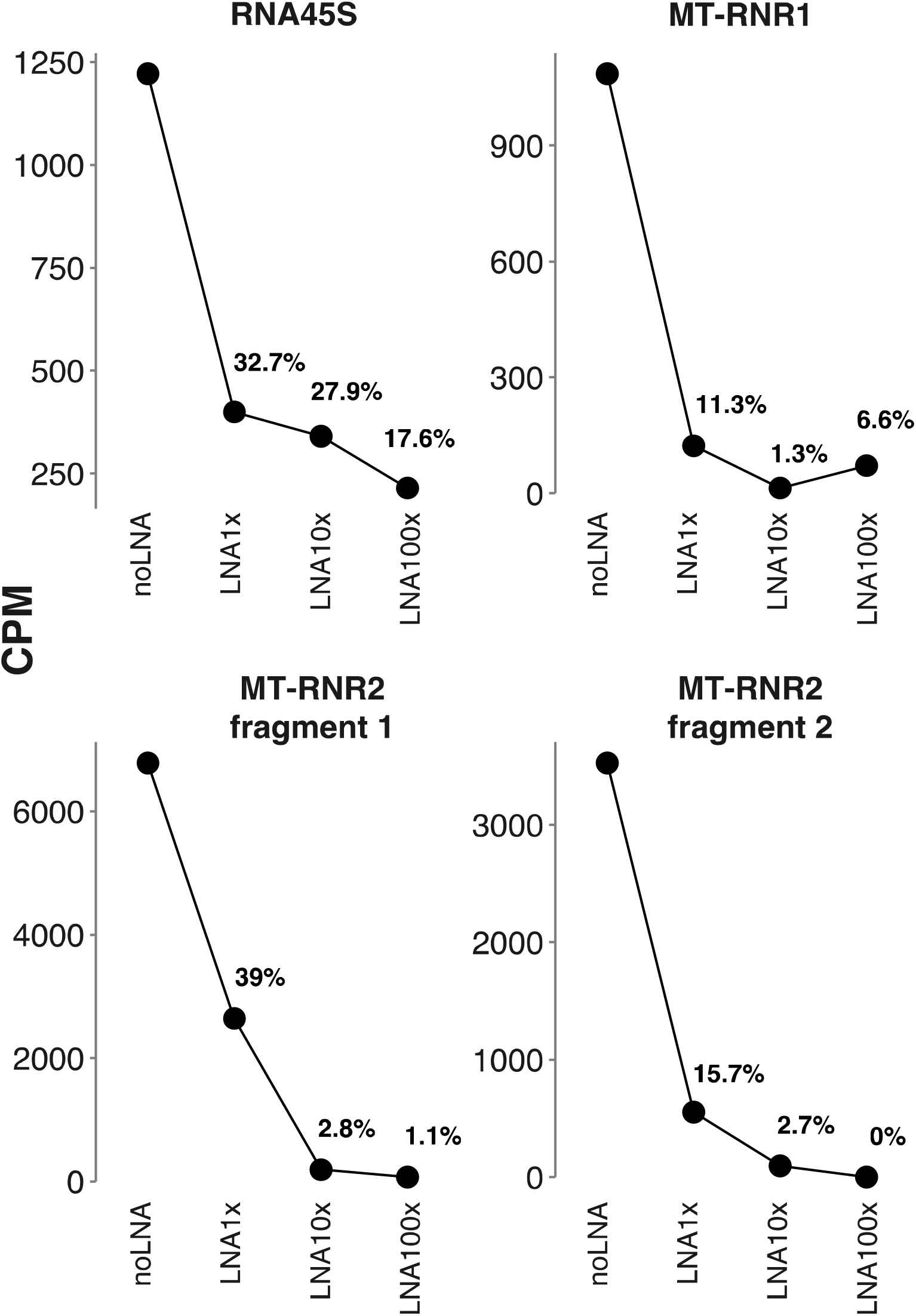
LNA oligonucleotide blocking of mtrRNA fragments (mitochondrially encoded 16S rRNA (two fragments), 45S pre-ribosomal RNA and mitochondrially encoded 12S rRNA) in Oxford Nanopore direct-cDNA sequencing. Counts per million (CPM) of the targeted transcripts for noLNA and LNA1x, LNA10x and LNA100x samples after Oxford Nanopore direct-cDNA sequencing. Fractions of CPM for LNA oligonucleotide treated samples relative to noLNA samples are shown as percentages.

### MALAT1 blocking in single-cell 3’ end sequencing of PBMCs

We finally evaluated if our method would also be applicable to single-cell RNA sequencing. More specifically, we designed two half-modified LNA oligonucleotides to block MALAT1 in single-cell 3’ end sequencing libraries of PBMCs. In PBMCs, MALAT1 can consume > 40% percent of reads through priming of internal poly(A) stretches (Supplemental Figure 6). The LNA oligonucleotides were added either before reverse transcription (pre-RT), which occurs in the gel bead-in-emulsion (GEMs), or before cDNA amplification, when the GEMs are pooled (pre-PCR). Both protocols show a decrease in MALAT1 reads (6-fold for the pre-RT and 4-fold for the pre-PCR blocked libraries) (Figure 6A). For some cell types, e.g., erythrocytes and regulatory T-cells, the initial MALAT1 proportions were higher, resulting in a more drastic reduction (Supplemental Figure 7). We observed a higher mitochondrial-derived RNA fraction for the pre-RT sample, which may indicate cell death (Figure 6A). The LNAs are, in this case, combined with living cells for 18 min. We therefore focused our analysis on the pre-PCR protocol. UMAP representation of cells based on single-cell RNA-seq data from both the pre-PCR blocking and standard protocol revealed tight clustering of cell types independent of protocol (Figure 6B), implying that the MALAT1 LNA oligonucleotide in the pre-PCR protocol has minimal impact on gene expression. This was further demonstrated by a perfect correlation (Spearman and Pearson correlation = 1.00) of gene expression values between the pre-PCR blocking and standard protocol (Supplemental Figure 8). We observed a significant increase, albeit with a small effect size, in the mean number of detected genes per cell in the pre-PCR protocol; 1173 genes with at least two counts in the control sample and 1192 in the pre-PCR blocking sample (p_t-test_ = 5.751e-05) (Figure 6A). The higher gene detection sensitivity might be related to the initial MALAT read fraction, the number of genes detected in the cells and the fraction of other highly abundant genes. The highest impact was seen in B memory cells where the mean number of detected genes increased from 1131 to 1235 (9.2% increase, padj, t-test = 0.026).

**Figure 6.**
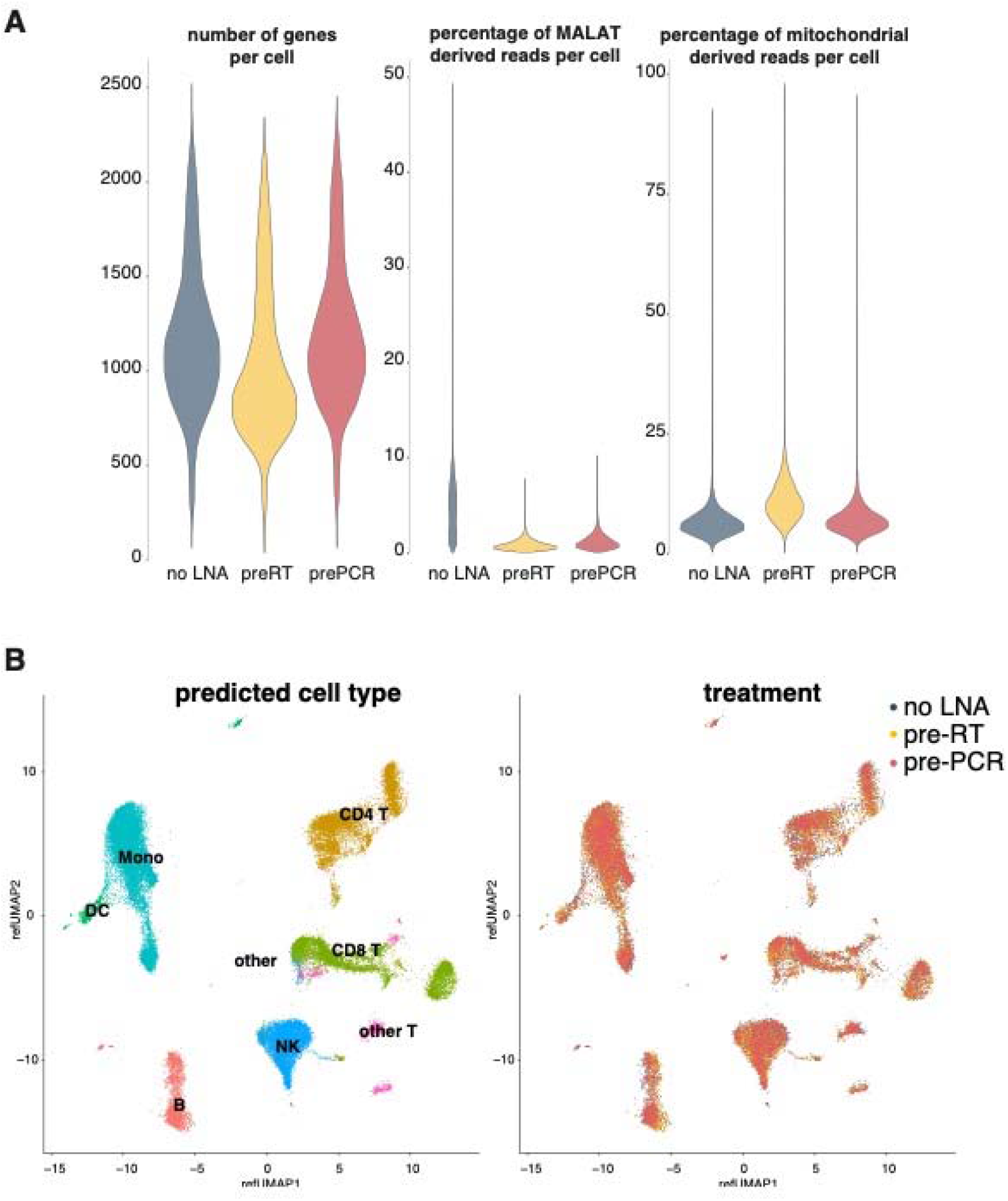
MALAT1 blocking in Chromium single-cell data. A. Quality metric distributions for the standard protocol, preRT and prePCR LNA oligonucleotide blocking, indicating a slightly higher number of detected genes in prePCR LNA oligonucleotide blocking, lower MALAT percentages in both preRT and prePCR LNA oligonucleotide blocking and higher mitochondrial RNA (a cell death proxy) in preRT LNA oligonucleotide blocking. B. Highly similar UMAP representations after integration of the samples are obtained for all protocols.

### LNA blocking simulation in whole blood 3’-end sequencing

Although our wet-lab experiments indicate that high-affinity binding oligonucleotide blocking can efficiently deplete transcripts of interest, it remains to be determined what the relationship is between initial abundance and level of depletion in order to offer substantial benefit (such as increased library complexity). Some of our applications had a more significant impact on the number of additionally detected genes and on coverage increase than others. We assume that this impact is highly dependent on the initial fraction of targeted reads, the depletion efficiency, and the sequencing saturation. In order to investigate in detail, we simulated different abundances and depletion efficiencies of beta-globin (HBB) using a publicly available whole blood 3’-end sequencing data, in which HBB accounted for 20.8% of all reads (Uellendahl-Werth et al., 2020)). Figure 7 shows how the number of detected genes (with at least 10 counts) increases linearly with increasing depletion efficiency at shallow sequencing depth but increases exponentially at higher sequencing depths. The linear relation for low sequencing depths probably results from unsaturated sequencing. The relation becomes more linear as the initial unwanted fraction lowers in higher sequencing depths. We conclude that even inefficient depletion of high-abundant transcripts provides a substantial gain in number of detected genes.

**Figure 7.**
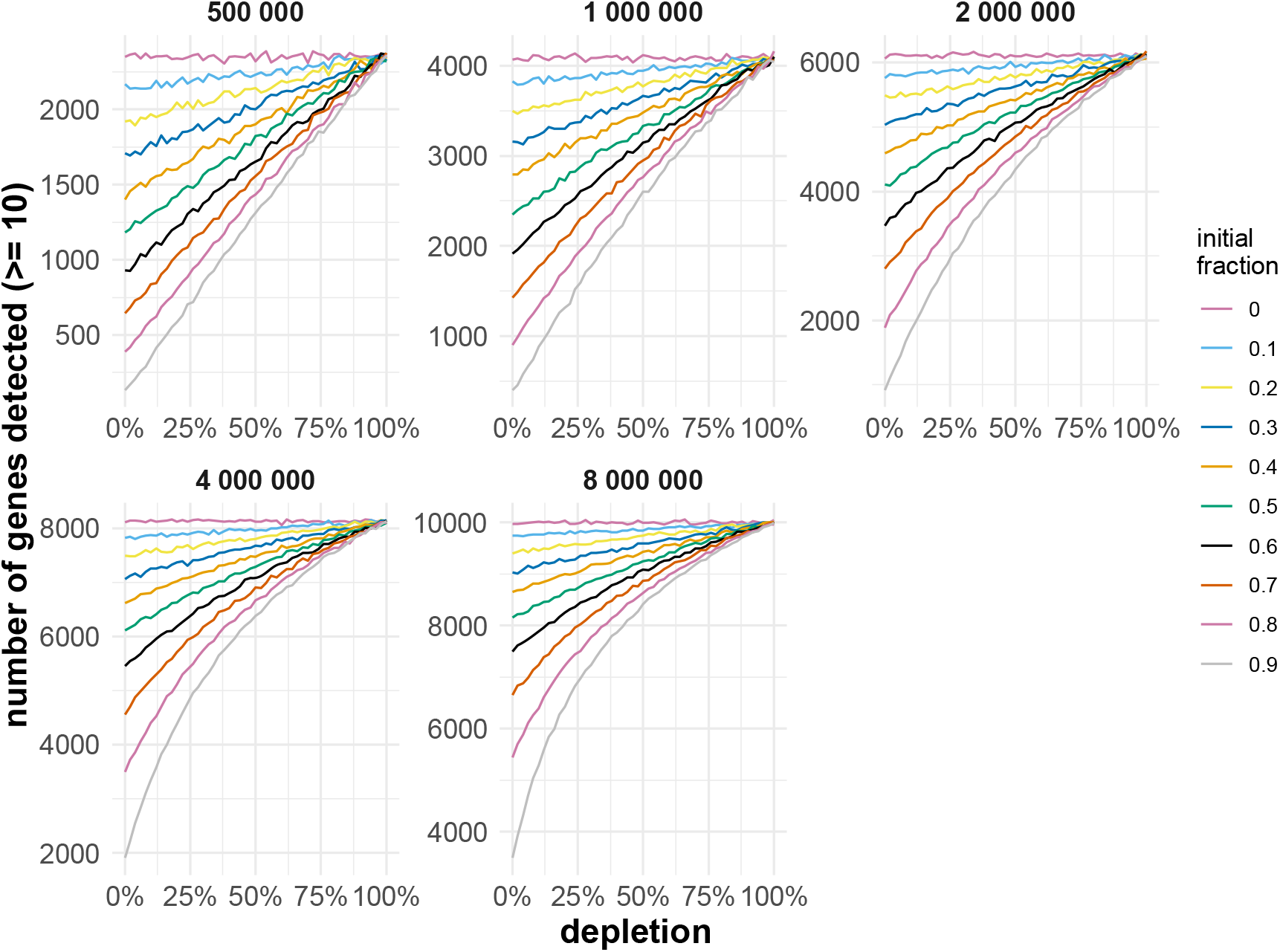
Simulations of HBB LNA oligonucleotide blocking in 3-end RNA sequencing data from whole blood. Each separate figure shows the number of genes with a count higher than 10 for each simulation with varying HBB depletion efficiency (0% means no depletion, 100% mean total depletion). The lines are colored by the initial fraction of HBB counts (of the total amount of counts). The graphs are separated by the number of counts the data was subsampled to.

## DISCUSSION

We demonstrate that high-affinity binding oligonucleotides can be applied to block reverse transcription and PCR amplification of various RNA transcripts in different RNA-seq library preparation protocols. We present a flexible and robust method that can drastically increase the detection and coverage of (low abundant) genes in the library. While LNA oligonucleotides have been used before to block PCR amplification (Hummelshoj et al., 2005), we provided evidence that such oligonucleotides can block both reverse transcription and PCR amplification, indicated by evident fragment depletion in the PCR-independent Oxford Nanopore Technologies protocol and the post-RT single-cell sample, respectively. Moreover, we demonstrate that the impact of blocking specific RNA fragments on gene detection and coverage strongly depends on the abundance of the blocked fragments, the transcriptome complexity and the sequencing depth. For example, the impact of RNY4 blocking on gene (i.e., miRNA) detection and coverage was much more significant for PRP than PFP, most likely because of higher RNY4 abundance in PRP. We expect a more pronounced impact in PFP samples at lower sequencing depths. We provide support for this hypothesis using simulated depletion experiments.

Our method has several advantages compared to existing protocols. First, the method only requires a single additional step that can be implemented in any RNA-seq library preparation workflow. Second, no nucleic acid sample or library material is lost because of enrichment or washing steps, which we believe has a positive impact on detection sensitivity, especially for low-input samples.

While we generally observe potent blocking of targeted transcripts, we also observe a few minor unwanted effects. First, in single cell RNA sequencing, adding LNA oligonucleotides to the GEMs during 3’ end sequencing resulted in a higher fraction of mtRNA reads. As living cells are incubated with LNA oligonucleotides for 18 min, the oligonucleotides may enter the cells and induce cell death. The larger the fraction of mtRNA for specific cell types, the fewer detected genes. Since adding the LNA oligonucleotides post-PCR also results in potent target blocking, we propose to use this approach instead. Second, the optimal concentration of LNA oligonucleotide may be application and target-dependent. A dedicated optimization step is warranted for optimal performance. This necessity is reflected in the single cell RNA sequencing experiment, where the benefit (in terms of the number of detected genes) depends on the cell type. Factors to consider are the original fraction of the targeted RNA transcript and the input RNA concentration of the library preparation protocol. We advise to combine samples in one single library prep to exclude batch effects, as is generally advised for RNA-seq experiments. Third, we observed a limited number of off-target effects upon adding specific LNA oligonucleotides (for instance, MT-AT8 and H4C3 in the 3’ end sequencing experiment). We did not observe significant sequence complementarity between the LNA oligonucleotides and the presumed off-targets. Nevertheless, off-target effects are not entirely unexpected given the relatively short length of the LNAs, their high RNA-binding capacity, and the small design space. The latter lowers the number of possible oligonucleotides and thereby the chances of designing one without off-target effects. Increasing oligonucleotide length or reducing the number of LNA nucleotides to lower binding affinity may improve specificity. A fourth limitation of the method is that it may only be applicable to small RNA sequencing or RNA-sequencing library prep methods employing an oligo(T) or a gene-specific RT primer. When the priming is random, it is impossible to design a single LNA oligonucleotide to block reverse transcription of the whole fragment. One option would be to design multiple LNA oligonucleotides spanning the entire transcript, but this could become prohibitively expensive, depending on the length of the fragment. Fifth, LNA synthesis is costly. Nevertheless, the amount of oligo that is required for efficient blocking is limited. Even at low synthesis scale, several hundreds of reactions can be performed, resulting in a limited per-sample cost. As fully and partially modified LNA oligonucleotides are equally efficient for YRNA depletion, partially modified LNA oligonucleotides could be used to further reduce oligo synthesis cost (although additional validation would be required as we only demonstrated this for a single RNA target sequence). Notably, blocking unwanted transcripts may also help reduce the sequencing cost. Finally, the observed shortening in read length with increasing LNA concentration in the Oxford Nanopore Technologies experiment is problematic, as it suggests off-target binding of the LNAs. Although the LNA oligonucleotides are expected to preferentially bind sequences with a lower number of mismatches, the steady decline in coverage towards the 5’-end of RNA transcripts points towards close to non-specific binding of the LNA oligonucleotides when supplied at high concentration. The possibility of the LNA oligonucleotides inhibiting the sequencing by binding to the final library can be dismissed by investigating the adaptor-to-adaptor reads (which signify complete sequencing of the read).

We believe the method presented here is versatile and can be used for other applications not investigated here, including hemoglobin mRNA blocking in whole blood samples (up to 70% of all mRNA in whole blood (Field et al., 2007)) or trypsin mRNA in pancreatic RNA samples. As we have shown, samples dominated by a few fragments have a higher potential of benefitting from LNA oligonucleotide-transcript blocking. We suggest the users to perform an initial computational analysis to define the expected benefit prior to implementing and optimizing our proposed method. While we only investigated mixtures of up to four different LNA oligonucleotides, it would be possible to combine more and block multiple fragments in one sample. Such mixtures can be designed specifically for unique and challenging sample types, containing several highly expressed, uninformative fragments (Hulstaert et al., 2020).

In conclusion, we present a novel and broadly applicable method to specifically block unwanted RNA transcripts during RNA sequencing library preparations by simply adding a target-specific high-afftinity oligonucleotide to the RT or PCR reaction.

## AVAILABILITY

The generated sequencing data is available through EGA with accession ID EGAS00001006023

## ACCESSION NUMBERS

EGAS00001006023

## SUPPLEMENTARY DATA

Supplementary Data statement:

Supplementary Data are available at NAR online.

## ACKNOWLEDGEMENT

**Celine Everaert:** Conceptualization, Methodology, Software, Formal Analysis, Resources, Data Curation, Writing – Original Draft, Writing – Review & Editing, Visualization, Funding Acquisition. **Jasper Verwilt:** Conceptualization, Methodology, Software, Formal Analysis, Investigation, Resources, Data Curation, Writing – Original Draft, Writing – Review & Editing, Visualization. **Kimberly Verniers:** Investigation, Writing – Review & Editing. **Niels Vandamme:** Investigation, Resources, Writing – Review & Editing, Funding Acquisition. **Alvaro Marcos Rubio:** Investigation, Resources, Writing – Review & Editing. **Jo Vandesompele:** Conceptualization, Resources, Writing – Review & Editing, Supervision, Funding Acquisition. **Pieter Mestdagh:** Conceptualization, Resources, Writing – Review & Editing, Supervision, Funding Acquisition.

