## Supplemental Materials for "Blocking abundant RNA transcripts by high-affinity oligonucleotides during transcriptome library preparation"

### Supplementary Files for *‘Blocking abundant RNA transcripts by high-affinity oligonucleotides during transcriptome library preparation’*

Celine Everaert^1,2^°, Jasper Verwilt^1,2^°, Kimberly Verniers^1,2^, Niels Vandamme^2,3^, Alvaro Marcos Rubio^1,2^, Jo Vandesompele^1,2#^ and Pieter Mestdagh ^1,2#^*

^1^ Department of Biomolecular Medicine, Ghent University, Ghent, Belgium

^2^ Cancer Research Institute Ghent, Ghent University, Ghent, Belgium

^4^ VIB Center for Inflammation Research, Vlaams Institute voor Biotechnologie, Zwijnaarde, Belgium

°"The authors wish it to be known that, in their opinion, the first two authors should be regarded as joint First Authors"

^#^”The authors wish it to be known that, in their opinion, the last two authors should be regarded as joint Last Authors”

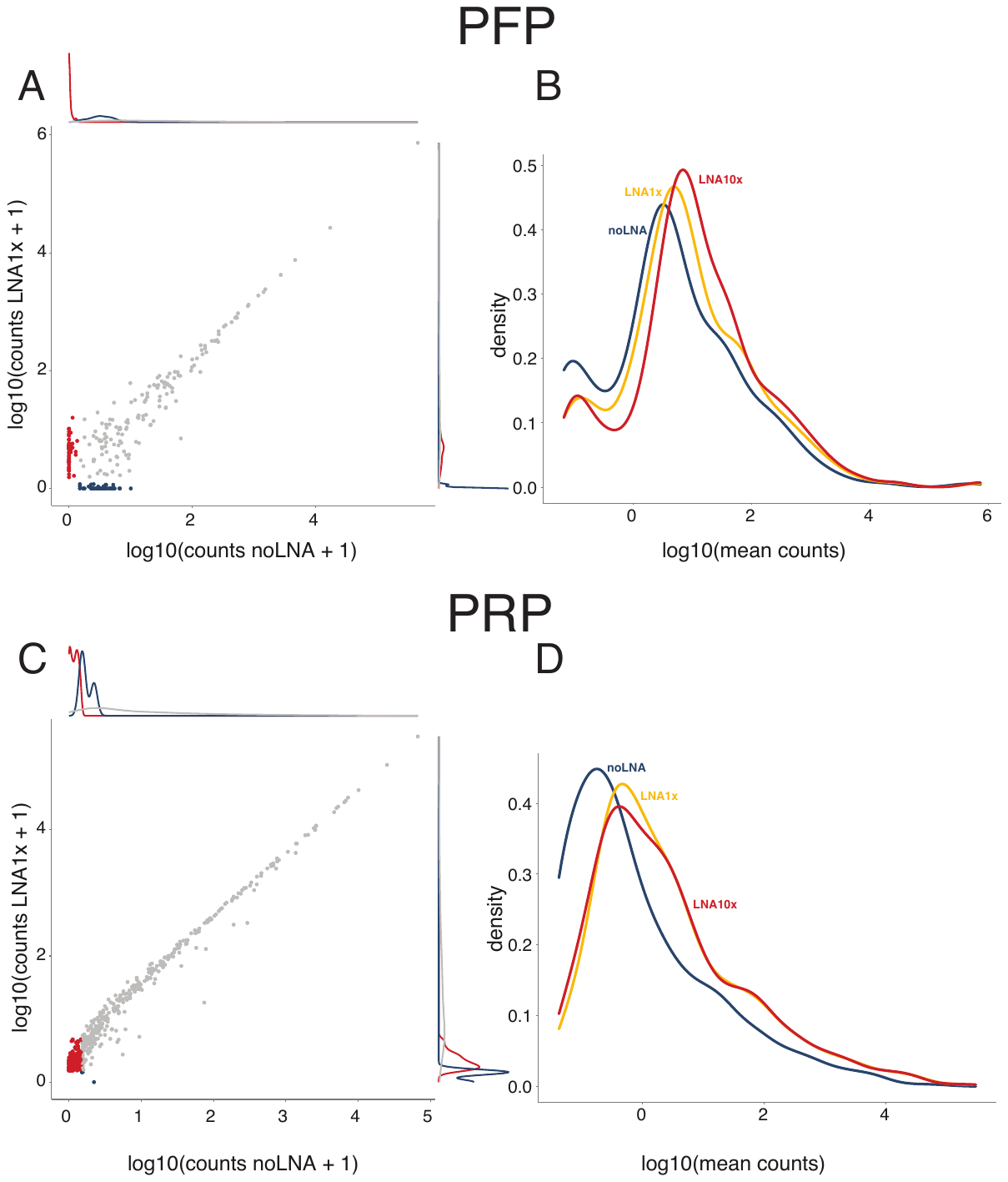
Supplemental Figure 1. **Count distributions of noLNA, LNA1x, and LNA10x in PFP and PRP.** A & C. Correlation plots are shown between the log_10_ of the counts (+1 to deal with 0’s) of the noLNA and LNA1x samples (averaged over technical replicates). Each dot represents a gene. Red dots are genes unique to LNA1x, blue dots are unique to noLNA. At each edge of the plot, distributions are shown of the values of the corresponding axis and treatment. A shows the results for PFF, C for PRP. B & D. Density plots of the log_10_ of the average counts (over technical replicates) for each of the sample. B shows the results for PFP, D for PRP.

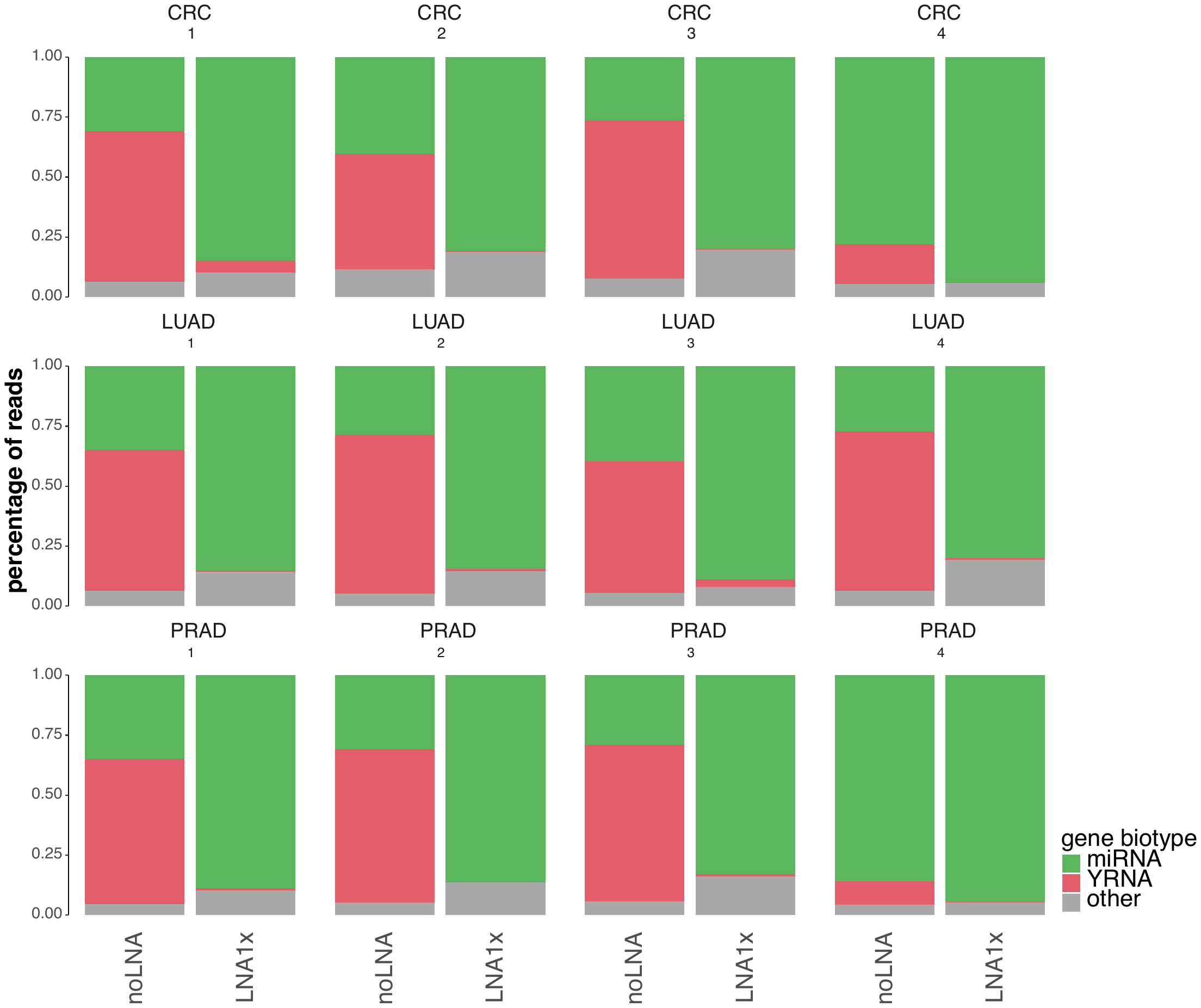
Supplemental Figure 2. **Read distribution for each sample and treatment.** The graphs are shown separately for each of the samples. For each graph, the percentage of reads going to either miRNA, YRNA, or other biotypes. The percentage of reads going to a certain biotype are shown and colored by biotype. (CRC = colorectal cancer, LUAD = lung adenocarcinoma, PRAD = prostate adenocarcinoma).

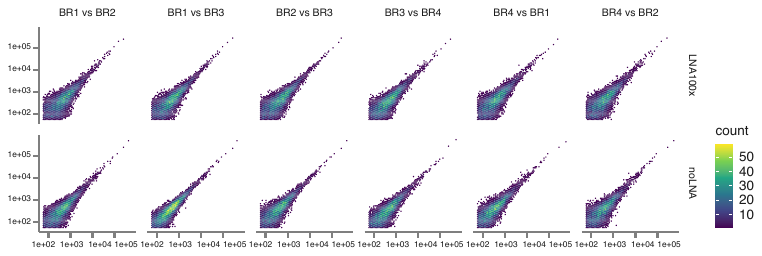
Supplemental Figure 3. **Correlation plots comparing all biological replicates.** Density plots for counts per million comparing each biological replicate. Each hexagon represents on the plot and they are colored by the number of genes falling within that region.

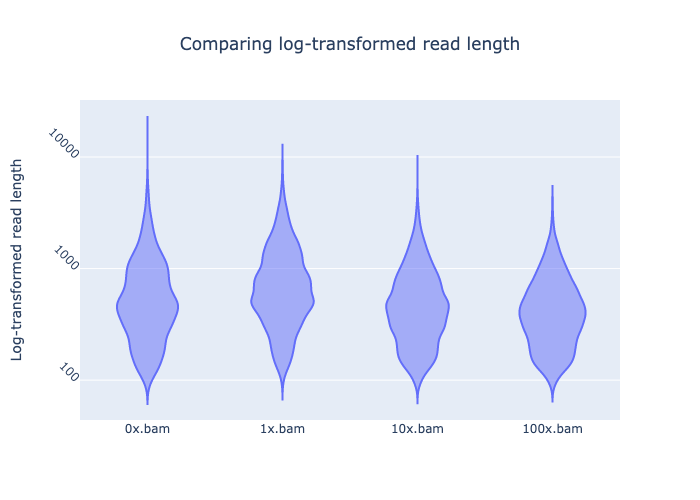

Supplemental Figure 4. **Length distribution of full-length transcripts after Oxford Nanopore direct-RNA sequencing.** For each of the samples (treated and non-treated) the distribution of the read lengths are visualized using violin plots. The y-axis is log10 transformed to allow for better investigation of the short reads. This plot is generated using NanoComp.

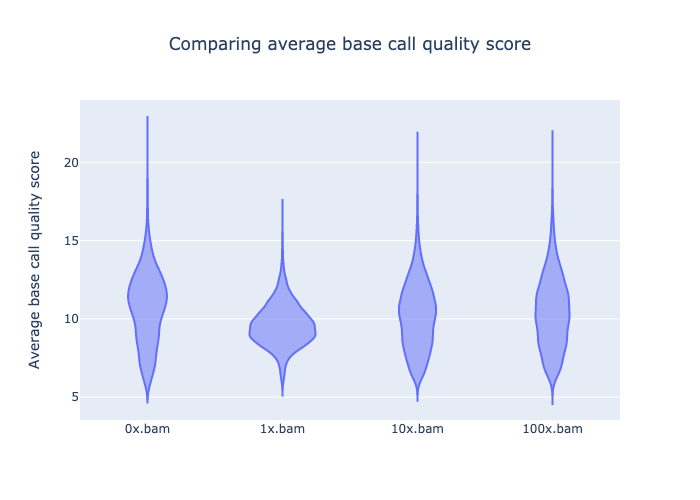

Supplemental Figure 5. **Quality score distribution of full-length transcripts after Oxford Nanopore direct-RNA sequencing.** For each of the samples (treated and non-treated) the distribution of the quality scores is visualized using violin plots. This plot is generated using NanoComp.

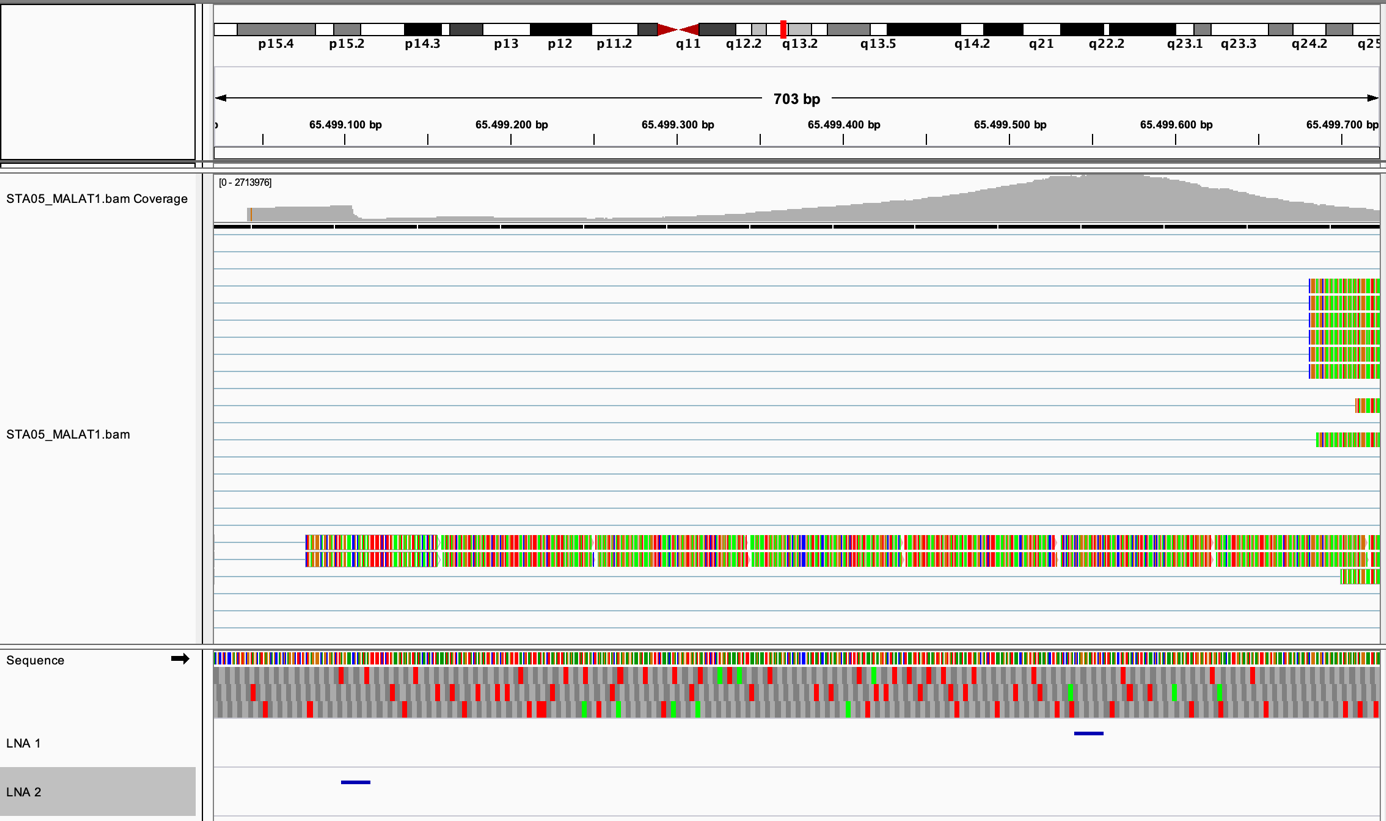

Supplemental Figure 6. **Illustration of problematic MALAT1 fragments and design space.** The upper track shows the reads mapping to MALAT1 for a single-cell 10X Chromium 3’ end sequencing library originating from PBMCs. The lower track shows where the designed LNA oligonucleotides are binding (two blue lines). This overview was created using IGV.

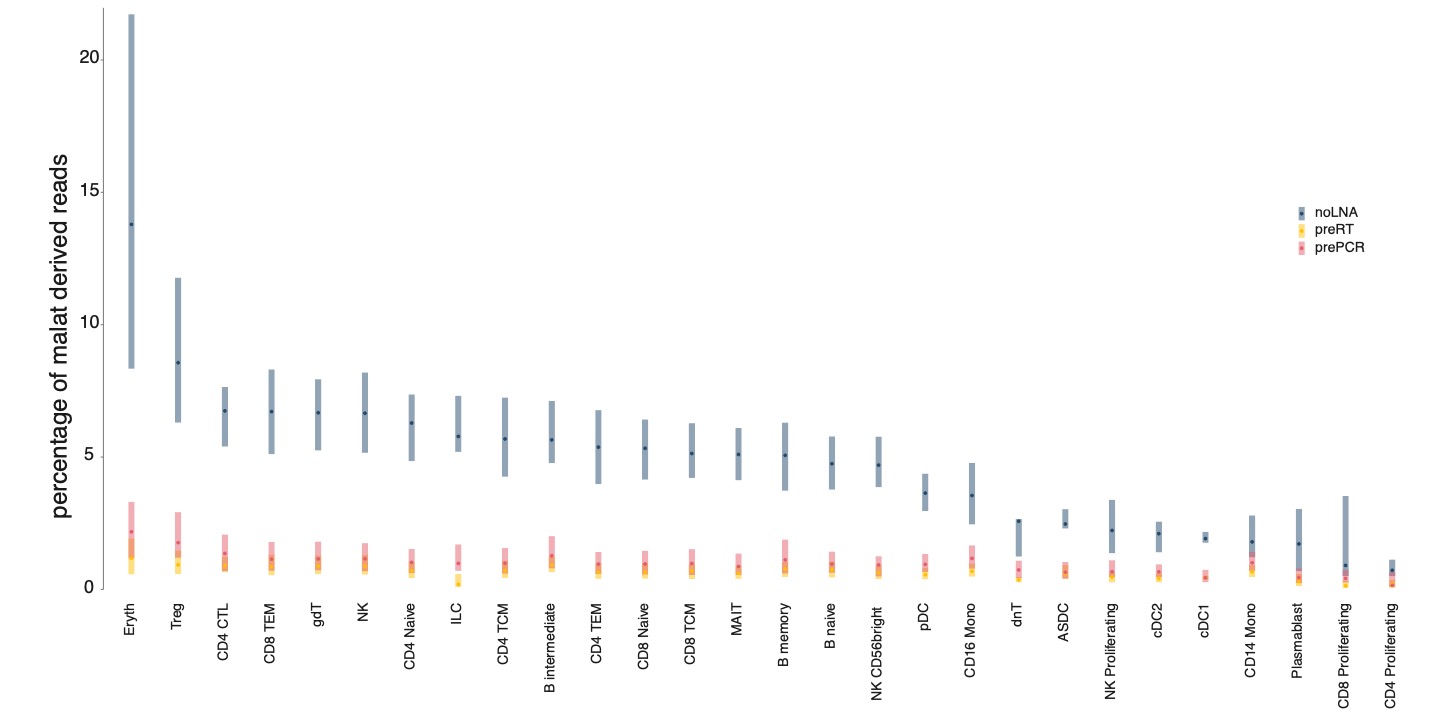

Supplemental Figure 7. **MALAT transcription in each cell type.** The percentage of reads mapping to MALAT are shown for each cell type. The averages and 95%-confidence intervals are shown and colored by the blocking regiment.

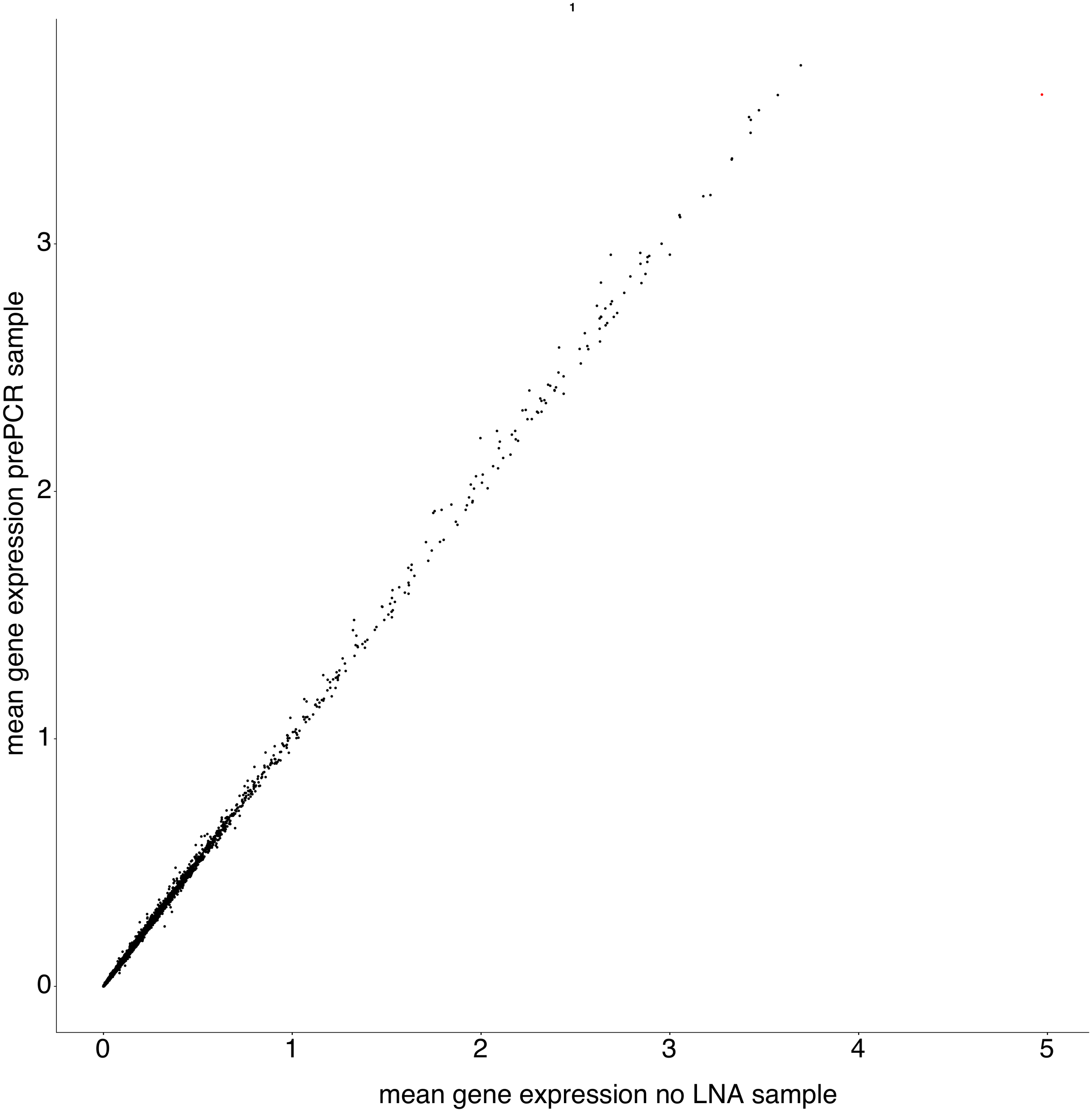

Supplemental Figure 8. **Correlation plot between prePCR and noLNA samples.** Counts are displayed as the median of the log_10_ of the counts+1. One is added to each count to deal with genes with 0 counts. Each dot is a gene.

Supplemental Table 1. **Sequences of designed synthetic oligonucleotides.** For each modified oligonucleotide the identification, gene target and sequence are provided. The sequences contain modification information in the generally accepted standard of notification.

| **oligo ID** | **targeted gene** | **sequence** |
| --- | --- | --- |
| RNY4_full_LNA | RNY4 | +A+C+C+C+A+C+T+A+C+C+A+T+C+G+G+A |
| RNY4_half_LNA | RNY4 | +AC+CC+AC+TAC+CA+TC+GG+A |
| RNY4_full_2'OMe | RNY4 | mA*mC*mC*mC*mA*mC*mT*mA*mC*mC*mA*mT*mC*mG*mG*mA |
| RNY4_half_2'OMe | RNY4 | mA*C*mC*C*mA*C*mT*A*C*mC*A*mT*C*mG*G*mA |
| RNY4_full_2’MOE | RNY4 | /52MOErA/*/i2MOErC/*/i2MOErC/*/i2MOErC/*/i2MOErA/*/i2MOErC/*/i2MOErT/*/i2MOErA/*/i2MOErC/*/i2MOErC/*/i2MOErA/*/i2MOErT/*/i2MOErC/*/i2MOErG/*/i2MOErG/*/32MOErA/ |
| RNY4_half_2’MOE | RNY4 | /52MOErA/*C*/i2MOErC/*C*/i2MOErA/*C*/i2MOErT/*A*C*/i2MOErC/*A*/i2MOErT/*C*/i2MOErG/*G*/32MOErA/ |
| RNA45S_full_LNA | RNA45S | +C+C+G+C+T+G+A+C+T+A+A+T+A+T+G+C |
| MT-RNR1_full_LNA | MT-RNR1 | +C+T+C+A+G+G +T+G+A+G+T+T +T+T+A+G |
| MT-RNR2_full_LNA_1 | MT-RNR2 | +A+G+A+C+G+G+G+T+G+T+G+C+T+C+T+T |
| MT-RNR2_full_LNA_2 | MT-RNR2 | +A+G+C+A+T+G+C+C+T+G+T+G +T+T+G+G |
| MALAT1_half_LNA_1 | MALAT1 | +GA+CG+CA+AT+TC+TC+CC+TG+C+G |
| MALAT1_half_LNA_2 | MALAT1 | +CT+TA+TC+TG+CG+GT+TT+CC+T+C |
